# A novel growth factor-dependent thermogenic brown adipocyte cell line from defined precursor cells

**DOI:** 10.1101/565168

**Authors:** Dagmar Kindler, Isabel S Sousa, Sabine Schweizer, Sarah Lerch, Martin Klingenspor, Stephan Herzig, Alexandros Vegiopoulos

## Abstract

Molecular pathways regulating brown adipocyte formation and metabolism can be exploited as targets for the treatment of obesity and disorders of glucose and lipid metabolism such as type-2 diabetes. Investigations in this direction require adequate cell models for brown adipocytes and their precursors. We report the establishment of a novel clonal cell line derived from defined Lin^−^Sca1^+^ adipocyte precursors from murine interscapular brown fat. In contrast to most currently available lines, immortalization was achieved by serial passaging without viral or genetic manipulation. Instead, the media were supplemented with basic fibroblast growth factor, which was required for the maintenance of stable long-term growth and immature morphology. BATkl2 cells differentiated to adipocytes with high efficiency upon standard adipogenic induction independently of PPARg agonists and even at higher passage numbers. BATkl2 adipocytes showed readily detectable Uncoupling protein 1 (Ucp1) protein expression and acutely responded to norepinephrine with increased Ucp1 mRNA expression, lipolysis and uncoupled mitochondrial respiration. Highly efficient siRNA-mediated knockdown was demonstrated in the growth state as well as in differentiating adipocytes, whereas plasmid DNA transfection was achieved in immature cells. These features make the BATkl2 cell line an attractive brown (pre)-adipocyte cell model.

## Introduction

Brown adipose tissue (BAT) is a major contributor to adaptive thermogenesis in mammals including humans and rodents. Its central function is the generation of heat through the combustion of chemical energy (Cannon *et al*. 2004, Diaz *et al*. 2014, Sidossis *et al*. 2015). In this way it can have a substantial contribution to organismal energy expenditure and thereby influence energy balance in the long-term. The amount and activity of BAT depends on environmental conditions, mainly temperature, as well as feeding status, age and obesity (Diaz *et al*. 2014, Li *et al*. 2014). Evidence from rodent models and human studies has established the protective function of BAT against obesity, the dysfunction of systemic glucose and lipid metabolism and associated diseases such as type-2 diabetes. Thus, enhancement of BAT function has become an actively pursued approach in the development of new therapies in this area (Betz *et al*. 2018, Moonen *et al*. 2019, Sidossis *et al*. 2015). In this direction it is critical to better understand the molecular regulation of BAT formation, maintenance and metabolism.

BAT develops mostly before birth and is located in several confined and species-specific anatomical sites (Diaz *et al*. 2014, Wang *et al*. 2016). The cell type responsible for heat generation is the brown adipocyte, characterized by multilocular lipid droplets, high mitochondrial content and high expression of the Uncoupling protein 1 (Ucp1). Although alternative mechanisms have been demonstrated, Ucp1-mediated energy dissipation plays a central role in heat generation (Cannon *et al*. 2004, Chouchani *et al*. 2019, Emont *et al*. 2019). This occurs through the uncoupling of the mitochondrial proton gradient and respiration from ATP synthesis. The metabolism of brown adipocytes is optimized for the efficient uptake and oxidation of substrates. This includes the release of intracellular fatty acids by lipolysis. Metabolic activation including lipolysis and mitochondrial uncoupling are controlled by various extracellular signals (Cannon *et al*. 2004, Emont *et al*. 2019). The major physiological stimulus is norepinephrine, which is locally released by the sympathetic nervous system.

Brown adipocytes are generated through the differentiation of immature mesenchymal precursor cells, also termed brown preadipocytes (Wang *et al*. 2016). This process involves lipid accumulation and mitochondrial biosynthesis and depends on the induction of a specific gene expression program including Ucp1 protein expression. Brown adipocyte formation is important beyond development and cell turnover, namely during the expansion of BAT upon prolonged cold adaptation. In this case, the proliferation of precursor cells contributes to increased capacity to form brown adipocytes (Lee *et al*. 2015, Nedergaard *et al*. 2019). BAT precursor cells with the capacity to form brown adipocytes have been identified by lineage tracing and flow cytometry in mice as Pdgfra^+^Sca1^+^ cells (Lee *et al*. 2015). We have shown that Lin(TER119/CD31/Cd45)^−^CD29^+^CD34^+^Sca1^+^ cells from murine interscapular brown fat efficiently form adipocytes *ex vivo* with a clearly distinct brown adipocyte expression profile compared to their counterparts from white adipose tissue (Bayindir *et al*. 2015, Rodeheffer *et al*. 2008).

The investigation of the molecular and biochemical mechanisms regulating precursor cell proliferation and differentiation as well as brown adipocyte metabolism requires appropriate cell culture models. Primary cells from the respective brown fat depots represent the most faithful approach. However, primary cells are only available in relatively low numbers and have limited capacity for proliferative expansion in culture. Human BAT is poorly accessible and only available from limited cohorts of patients. Primary cells can be functionally expanded for a limited number of passages upon viral transduction with oncogenes. Furthermore, several murine and human cell lines have been established through immortalization by stable expression of viral oncogenes or telomerase subunits (Table 1) (Cannon *et al*. 2001, Hirschberg *et al*. 2011, Klein *et al*. 1999). These preadipocytes can be induced to differentiate to brown adipocytes in culture. However, the expression of oncogenes and cell cycle regulators can confound conclusions on the regulation of precursor proliferation and can interfere with differentiation. In addition, currently available cell lines can have limitations such as the requirement for certain inducers for differentiation, limited capacity for differentiation in the long-term or partial lack of key brown adipocyte features.

**Table 1.**
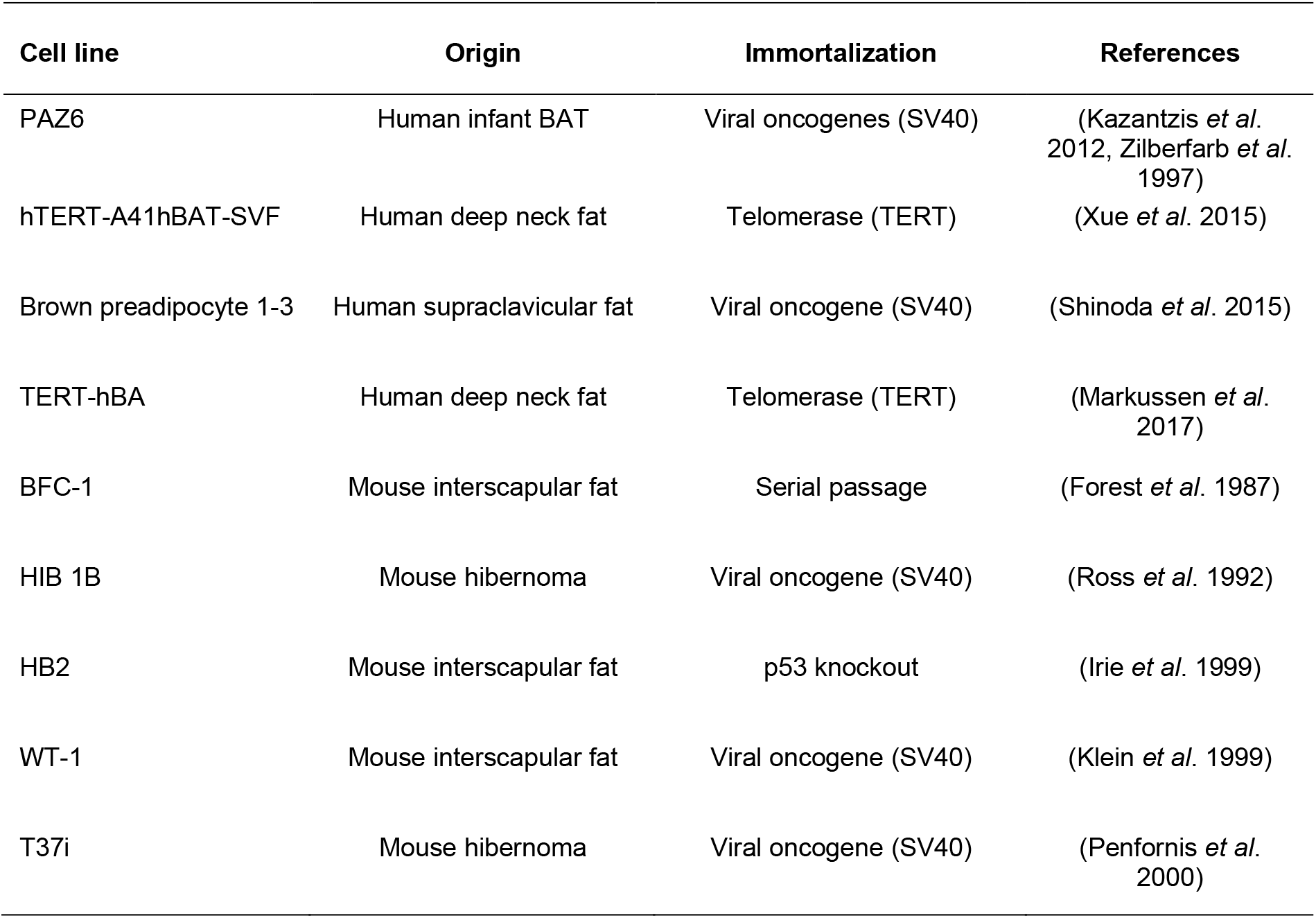
Overview of established murine and human brown preadipocyte cell lines.

We sought to establish a stable brown adipocyte precursor cell line from a defined primary cell population and without the use of viral/genetic manipulation. To this end we applied the 3T3 immortalization method with the addition of a physiological growth factor on primary precursor cells isolated from murine interscapular brown fat.

## Results

### Establishment of a stably growing bFGF-dependent clone from interscapular brown adipose tissue Lin^−^Sca1^+^ precursor cells

With the aim of obtaining a defined suitable cell population for the immortalization of brown adipocyte precursor cells, we isolated Lin^−^Sca1^+^ cells from murine interscapular brown adipose tissue by magnetic separation. As described previously, this population represents a good approximation of the Lin^−^CD29^+^CD34^+^Sca1^+^ precursor/progenitor population (*Babaei et al. 2018*, Bayindir *et al*. 2015). To apply the 3T3 immortalization approach we serially passaged the cells over several weeks (Wu *et al*. 2012). However, we included basic fibroblast growth factor (bFGF) in the media, which has been shown to promote mesenchymal/fibroblastic cell proliferation and the maintenance of adipogenic differentiation capacity (Hebert *et al*. 2009, Marquez *et al*. 2017, Widberg *et al*. 2009). The cultures reached stable exponential growth by approximately 15 passages and cloning was performed by serial dilution at 31 passages (Figure 1A). The clone designated BATkl2 showed stable exponential growth in the presence of bFGF, which was maintained for over 25 passages after a freeze-thaw cycle (Figure 1B). Notably, BATkl2 displayed poor adherence to the standard tissue culture coating, resulting in loss of cells. Therefore, the cells were transferred and permanently grown on gelatin-coated dishes, which resulted in improved plastic adhesion, reduced intercellular adhesion and increased growth, at least in the presence of bFGF (Figure 1C-I). The continuous culture with bFGF was associated with a denser, more immature cell morphology and higher growth rate (Figure 1D,F,H-J).

**Figure 1.**
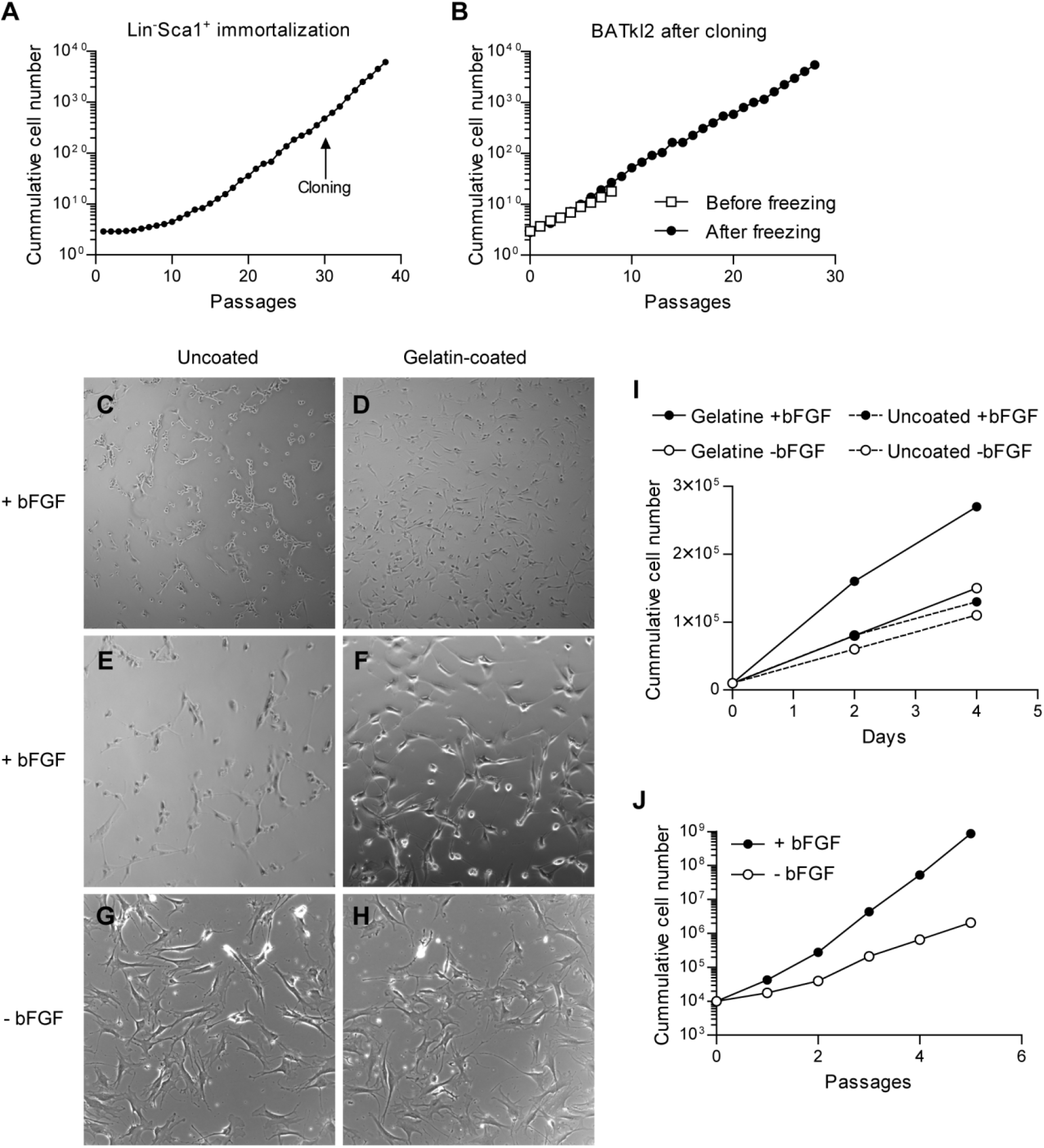
Establishment of a stably growing bFGF-dependent clone from interscapular brown adipose tissue Lin^−^Sca1^+^ precursor cells. (A) Growth curve of Lin^−^(Ter119/CD31/CD45)Sca1^+^ cells from interscapular brown adipose tissue during the 3T3 immortalization process in the presence of bFGF. “Cloning” indicates the passage at which clones were isolated by limiting dilution. “Passages” indicates the number of passages after plating out the primary cells. (B) Growth curve of the clone BATkl2, before and after cryopreservation. “Passages” indicates the number of passages after establishment of the clone or after freeze/thaw. (C-H) Phase-contrast images of BATkl2 cultures with or without gelatin coating and bFGF as indicated. 50x magnification (C,D); 100x magnification (E-H). (I) Growth curve of clone BATkl2 with or without gelatin coating and bFGF as indicated. (J) Growth curve of clone BATkl2 on gelatin-coated wells with or without bFGF. “Passages” indicates the number of passages from the beginning of the experiment shown.

### Efficient brown adipogenic differentiation of BATkl2 cells

BATkl2 cultures were induced to differentiate with a standard adipogenic cocktail on gelatin-coated dishes. Widespread adipogenic differentiation could be observed at 8 days after induction (Figure 2A,C,D). Differentiation failed in the absence of gelatin coating due to poor adhesion and loss of differentiating adipocytes (data not shown). Uncoupling protein 1 (Ucp1) is the prime expression marker for brown adipocytes and key mediator of the mitochondrial uncoupling process. Differentiated BATkl2 cells had 1000-fold higher *Ucp1* mRNA expression levels compared to non-induced BATkl2 cultures (Figure 2G). *Ucp1* expression could be further elevated by acute treatment with norepinephrine (Figure 2G). Importantly, Ucp1 protein could be robustly detected by Western blotting in differentiated but not in undifferentiated BATkl2 cultures (Figure 2H). Addition of rosiglitazone during the first 2 days of differentiation resulted in slightly increased adipogenic differentiation and potentiated *Ucp1* expression by more than 10-fold (Figure 2A-G). However, *Ucp1* mRNA levels were not sensitive to norepinephrine treatment in cultures induced with rosiglitazone. Irrespective of rosiglitazone treatment, the differentiation capacity and norepinephrine responsiveness were maintained for at least 10-15 passages after replating of cryopreserved cells, when passaging at the recommended low densities (data not shown).

**Figure 2.**
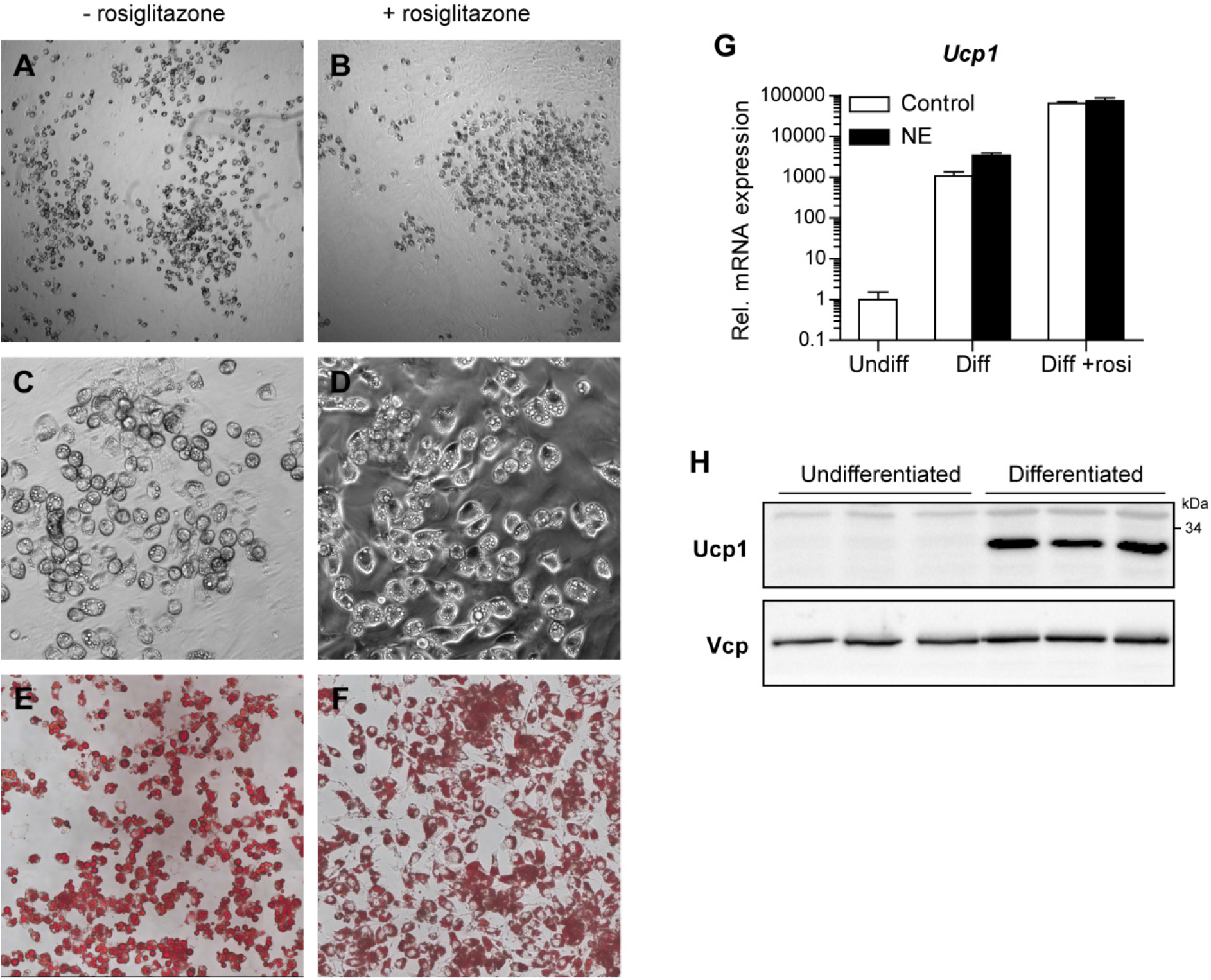
Efficient brown adipogenic differentiation of BATkl2 cells. BATkl2 cells (after cryopreservation) were differentiated for 8 days on gelatin-coated wells with or without rosiglitazone (rosi) during the 2-day induction phase. (A-D) Phase-contrast images at 50x (A,B) or 200x (C,D) magnification. (E,F) Microscopy images (100x magnification) of cultures stained with Oil Red O. (G) *Ucp1* mRNA expression, determined by qRT-PCR following 3-hour terminal treatment with norerpinephrine (NE, 0.5 μM) (n=3 cultures). Data represent mean and standard deviation. Undiff: undifferentiated; Diff: differentiated. (H) Ucp1 protein expression before and after differentiation, as determined by Western blot with Valosin containing protein (Vcp) as loading control (n=3 cultures).

### Norepinephrine-dependent lipolysis and uncoupled mitochondrial respiration in BATkl2 adipocytes

To determine the capacity of BATkl2 differentiated adipocytes to respond metabolically to norepinephrine and beta-adrenergic stimulation, we measured the lipolysis-dependent release of non-esterified fatty acids (NEFA). Treatment of differentiated BATkl2 cultures with 0.5 μM norepinephrine for 3 hours caused an approximately 10-fold increase in the concentration of NEFA in the culture supernatants (Figure 3A), implying a potent activation of triglyceride lipolysis. This effect could be recapitulated by treatment with 10 μM of the β-adrenergic agonist isoproterenol (Figure 3A). Furthermore, we asked whether BATkl2 adipocytes were able to activate uncoupled mitochondrial respiration in response to norepinephrine. To this end we differentiated BATkl2 cells in gelatin-treated XF96 microplates and measured their oxygen consumption rate (Seahorse analysis). BSA was included in the media to reduce non-specific effects of lipolysis-derived NEFAs on mitochondrial respiration (Li *et al*. 2014). Treatment with oligomycin inhibits ATP synthase and reveals the rate of uncoupled respiration. Stimulation with norepinephrine in the presence of oligomycin acutely increased oxygen consumption to a substantial proportion of the maximal respiratory capacity, determined by treatment with the general uncoupler FCCP (Figure 3B).The data suggest that BATkl2 differentiated adipocytes respond to norepinephrine by activating lipolysis and uncoupled mitochondrial respiration, representing central metabolic functions of brown adipocytes.

**Figure 3.**
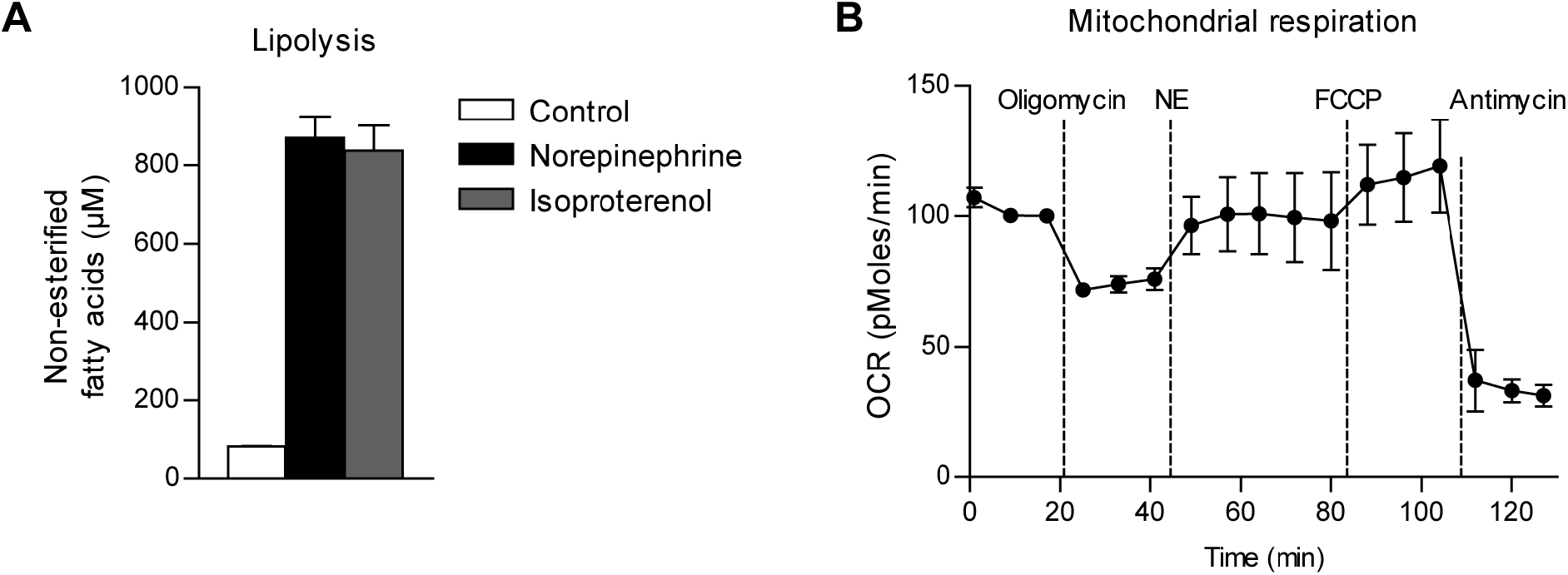
Norepinephrine-dependent lipolysis and uncoupled mitochondrial respiration in BATkl2 adipocytes. (A) Non-esterified fatty acid concentration in the supernatant of BATkl2 cultures following differentiation and terminal 3-hour treatment with the indicated β-adrenergic agonists (norepinephrine at 0.5 μM, isoproterenol at 10 μM; n=3 cultures). (B) Mitochondrial respiration of BATkl2 cultures following differentiation, as determined by Seahorse Extracellular Flux analysis in the presence of 2% bovine serum albumin. The vertical dashed lines represent the injection of the indicated substances (n=12). NE: norepinephrine 0.5 μM; FCCP: trifluorocarbonylcyanide phenylhydrazone; OCR: oxygen consumption rate. Data represent mean and standard deviation.

### Transfection efficiency in BATkl2 cells: siRNA and plasmid-mediated expression

We next tested whether BATkl2 cells were amenable to RNA interference using simple siRNA transfection. Knockdown experiments in the undifferentiated state are relevant for the investigation of gene function in precursor cell proliferation and early differentiation processes. BATkl2 cells were transfected under general growth conditions with siRNA targeting the *Prkaca* mRNA or with non-targeting siRNA (NC) and analyzed 2 days later. Transfection with siNC caused an unspecific 2-fold increase in the expression of *Prkaca* mRNA compared to non-transfected cells (Figure 4A). However, siPrkaca-transfected cells displayed a 20-fold reduction in *Prkaca* expression compared to siNC, corresponding to more than 90% knockdown efficiency (Figure 4A). The investigation of gene function in adipocytes independently of any effects on precursor proliferation and differentiation requires transfection after commitment to differentiation. To this end we transfected BATkl2 cells 5 days after the induction of differentiation with siPrkaca which resulted in approx. 90% knockdown of *Prkaca* mRNA compared to siNC (Figure 4B). Finally, we tested the possibility of transfecting BATkl2 cells with plasmid DNA in the general growing mode, aiming at the overexpression of genes of interest. GFP-expressing cells were readily detectable by fluorescence microscopy 24 hours after transfection with a GFP-encoding expression vector (Figure 4C,D). The transfection efficiency was of intermediate level. In conclusion, BATkl2 cells were highly amenable to siRNA-mediated gene knockdown in the undifferentiated as well as differentiated states and can in principle be transfected with DNA expression vectors.

**Figure 4.**
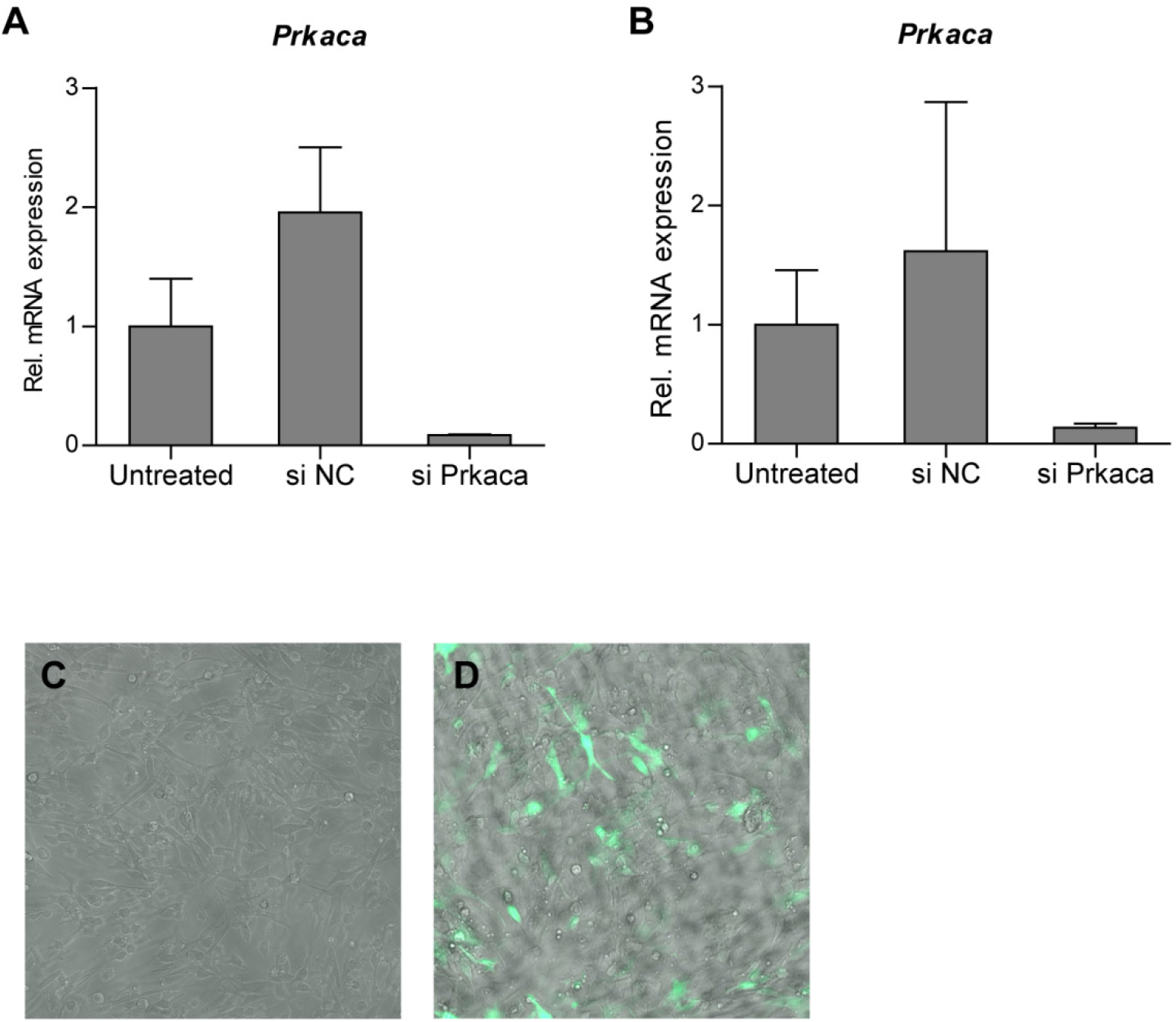
Transfection efficiency in BATkl2 cells: siRNA and plasmid-mediated expression. (A) *Prkaca* mRNA expression of BATkl2 cells 2 days after transfection with the indicated siRNA in the undifferentiated state (n=3 cultures). si NC: non-specific siRNA control. (B) *Prkaca* mRNA expression of BATkl2 cells at day 8 of differentiation after transfection with the indicated siRNA at 5 days of differentiation (n=3 cultures). (C,D) Merged phase-contrast and fluorescence microscopy images of BATkl2 cells 24 hours after transfection with pEGFP-N1 plasmid in the undifferentiated state. (C) non-transfected cultures. Data represent mean and standard deviation.

## Discussion

The novel BATkl2 cell line features a number of unique advantages compared to established human and murine brown preadipocyte cell lines. (1) To our knowledge it is the first BAT-derived cell line with defined cell type of origin, namely Lin^−^Sca1^+^ cells representing precursor cells. This can be of importance when it comes to investigations relevant to the currently ill-defined cellular differentiation hierarchies in adipose tissue. (2) The key distinguishing feature is that the immortalization process of BATkl2 cells is not dependent on viral oncogenes or other exogenous cell cycle regulator genes. Most currently available cell lines and immortalization protocols rely on such genes (Table 1) and this is a disadvantage given the ability of oncogenes to directly and constitutively influence cell proliferation and adipogenic differentiation. For instance, SV40 oncogenes have been shown to inhibit adipogenesis (Cherington *et al*. 1988). The proliferation rate and immature cell morphology of BATkl2 cells depended on continuous culture in the presence of the growth factor bFGF even after more than 40 passages post isolation. Under these conditions BATkl2 showed remarkably stable growth and maintained differentiation capacity at high passage number and after cryopreservation. (3) BATkl2 showed high differentiation efficiency including Ucp1 protein expression without the need of treatment with agonists of the Peroxisome Proliferator-Activated Receptor gamma (PPARγ). For instance, treatment with rosiglitazone or indomethacin is a requirement for many established BAT cell lines and interferes with central adipogenic pathways, potentially reducing the sensitivity of certain assays. (4) BATkl2-derived adipocytes displayed multiple key brown adipocyte features, in particular easily detectably Ucp1 protein expression as well as the acute induction of gene expression, lipolysis and uncoupled mitochondrial respiration in response to norepinephrine stimulation. (5) BATkl2 cells were amendable to siRNA transfection and showed highly efficient gene knockdown in the growth state as well as in differentiating adipocytes without the need of cell detachment (Isidor *et al*. 2016). In addition, transfection of undifferentiated BATkl2 cells with plasmid DNA reached an efficiency which is probably sufficient for reporter assays. One disadvantage of the BATkl2 cell line is the requirement for gelatin coating of the cell culture surface to achieve stable adhesion, which we speculate to be related to the high propensity for adipogenic differentiation and the associated rounding of cell shape.

Taken together, the BATkl2 cell line represents a novel attractive model for the investigation of brown adipocyte differentiation and thermogenic metabolism. It is suitable for experiments requiring extensive culture expansion and has the potential for use in high-throughput cell-based assays. The characteristics of the BATkl2 cell line may result in improved validity of conclusions related particularly to the regulation of precursor cell proliferation and differentiation. Finally, the growth factor-dependent immortalization approach presented here may facilitate the non-viral establishment of novel cell lines from murine and human brown adipose tissue.

## Materials and Methods

### Tissue dissociation and cell isolation

Female NMRI mice (Charles River WIGA GmbH, Sulzfeld, Germany) were housed at ambient temperature with 12-hour light-dark cycle on chow (Kliba Nafag #3437, Provimi Kliba, Kaiseraugst, Switzerland). Animal handling and experimentation was performed in accordance with the European Union directives and the German animal welfare act (Tierschutzgesetz) and approved by local authorities (Regierungspräsidium Karlsruhe). Interscapular brown fat was dissected from 6 week-old mice and cleared of surrounding white-appearing fat tissue. A single cell suspension of the stromal-vascular fraction was obtained by enzymatic tissue dissociation as previously described (Bayindir *et al*. 2015). Lin^−^ Sca1^+^ cells were isolated by the MACS cell separation procedure described in Bayindir *et al*. (Bayindir *et al*. 2015). Briefly, following magnetic depletion of Lin^+^ cells with antibodies against Ter119 (clone TER-119), CD31 (clone 390) and CD45 (clone 30-F11) (ebioscience), Sca1^+^ cells were isolated with anti-Sca1-PE-Cy7 (D7, ebioscience) and anti-Cy7 MicroBeads (Miltenyi Biotec, 130-091-652).

### Immortalization, clone derivation and maintenance culture

Freshly isolated Lin^−^Sca1^+^ cells (2.5*10^4^ per cm^2^) were plated out initially on standard tissue culture-grade plates in DMEM (high glucose) supplemented with 10% fetal calf serum (FCS), penicillin/streptomycin (Life Technologies) and 10 ng/ml recombinant murine bFGF (R&D Systems). Cells were passaged every 3-6 days according to current growth rate whereby 5*10^3^-2.5*10^4^ cells per cm^2^ were seeded. Clonal derivation of BATkl2 was performed at passage 31 (from primary cell isolation) by limiting dilution in 96-well plates. Single cell cultures were confirmed microscopically. For further culturing, BATkl2 was plated at 1-1.5*10^3^ cells per cm^2^ on gelatin-coated tissue culture plates in DMEM, 10% FCS, pen/strep and 10 ng/ml bFGF. Gelatin coating was performed by incubation of the surfaces with 0.2% autoclaved gelatin (from bovine skin, Sigma G9391) at 37°C for 15 minutes. Cultures were maintained at relatively low density, media were replaced every 2-3 days and passaging was performed by trypsinization.

### Adipogenic differentiation

Cells were plated on gelatin-coated wells at 1-3*10^4^ cells per cm^2^ in growth medium. At 90-100% confluence, media were replaced with DMEM (high glucose), 10% FCS, Pen/Strep (Life Technologies), 500 nM dexamethasone, 500 μM 3-isobutyl-1-methylxanthine (IBMX), 1 μg/ml recombinant human insulin, 3 nM triiodothyronine (T3) (Sigma) with or without 1 μM rosiglitazone (Biomol, Germany). The following day media were replaced by fresh. 2 days after differentiation induction, media were replaced with DMEM (high glucose), 5% FCS, Pen/Strep, 1 μg/ml recombinant human insulin and 3 nM triiodothyronine (T3). Media were replaced by fresh every 2 days. 6-8 days after differentiation induction, media were replaced by DMEM (high glucose), 5% FCS and Pen/Strep. Cells were harvested or analyzed at 8-10 days after differentiation induction. Wherever indicated, norepinephrine (Sigma) was added at 500 nM.

### siRNA transfection

FlexiTube siRNA Mm_Prkaca_1 (SI01388331, QIAGEN, Germany) and AllStars Negative Control siRNA (QIAGEN) were used. For transfection of undifferentiated cells, cells were seeded in 24-wells coated with 0.2% gelatin at 2-4*10^4^ cells per well in DMEM, 10% FCS and 10 ng/ml bFGF one day before transfection. For transfection of cells after the induction of differentiation, cells were induced to differentiate for the indicated time and media were replaced with DMEM (high glucose), 5% FCS, 1 μg/ml recombinant human insulin and 3 nM triiodothyronine (T3) one day before transfection. The transfection mix was prepared by combining a dilution of 2 μl RNAiMax (Life Technologies) in 14 μl OptiMEM (Life Technologies) with an appropriate dilution of siRNA in 16 μl OptiMEM (for one 24-well). Following 5 min incubation at room temperature, the transfection mix was diluted with the corresponding culture media to 500 μl and used to replace the current culture media. The siRNA concentration in the culture media was 20 nM. Culture media were replaced by fresh the following day.

### Plasmid DNA transfection

For plasmid DNA transfection of undifferentiated cells, cells were plated at 4*10^4^ cells per 24-well the day before transfection. 400 ng of pEGFP-N1 (Clontech) and 1.5 μl Attractene Transfection Reagent (QIAGEN) were diluted into 60 μl DMEM and incubated for 10 minutes at room temperature. The transfection mix was diluted to 500 μl DMEM supplemented with 10% FCS and used to replace the media of one 24-well.

### Oil Red O staining

Oil Red O staining was performed as previously described (Babaei *et al*. 2018).

### Lipolysis assay

Differentiated cells were incubated with Krebs-Ringer buffer for 2 hours before treatment with 0.5 μM norepinephrine or 10 μM isoproterenol (Sigma) in Krebs-Ringer buffer supplemented with 5% bovine serum albumin (BSA), 5 mM D-glucose and 25 mM HEPES/KOH pH 7.4 for 3 hours. Non-esterified fatty acid content in the culture supernatants was determined using the colorimetric NEFA-HR(2) assay kit (Wako Chemicals) according to manufacturer’s instructions.

### Mitochondrial respiration assay

Mitochondrial respiration was measured in live cells using the XF96 Extracellular Flux Analyzer (Seahorse Bioscience, Copenhagen, Denmark) as previously described (Li *et al*. 2014). Briefly, BATkl2 cells were plated on gelatin-coated XF96 microplates and treated for adipogenic differentiation as described above (with rosiglitazone for 2 days). The measurement was performed on day 8 of differentiation. Cells were treated sequentially at the indicated time points with 0.5 μM oligomycin, 0.5 μM norepinephrine, 1 μM trifluorocarbonylcyanide phenylhydrazone (FCCP) and 5 μM antimycin A in the presence of 2% BSA. The oxygen consumption rate was calculated with XF-96 software.

### qRT-PCR mRNA expression analysis

Total RNA was isolated from cultured BATkl2 cells using Qiazol (QIAGEN) according to the manufacturer’s instructions. cDNA synthesis was performed using the First Strand cDNA Synthesis Kit (Fermentas/Thermo Scientific). Quantitative RT-PCR was performed on a StepOnePlus Real Time PCR System (Life Technologies) using Taqman Gene Expression Assays for *Ucp1* and *Tbp* (Life Technologies) or Mm_Prkaca_1_SG and Mm_Tbp_1_SG QuantiTect Primer Assay with SYBR Green for *Prkaca* and *Tbp* respectively (QIAGEN). Relative expression values were calculated using the ΔΔCt method including normalization with the *Tbp* values.

### Western blotting

Western blotting was performed as previously described (Bayindir-Buchhalter *et al*. 2018) with anti-Ucp1 (1:1000; PA1-24894; ThermoFisher Scientific) and anti-Vcp (1:10000; ab11433; Abcam) in TBS with 0.1% Tween20 and 1% BSA.

## Acknowledgements

We’d like to apologize to any scientists having contributed to the generation of BAT-derived cell lines that were not cited or mentioned here. This work was supported by the Deutsche Forschungsgemeinschaft (HE 3260/8-1) and the Helmholtz Association (“Metabolic Dysfunction”). The authors have no conflict of interest to declare.

## References

Babaei R, Schuster M, Meln I, Lerch S, Ghandour RA, Pisani DF, Bayindir-Buchhalter I, Marx J, Wu S, Schoiswohl G, Billeter AT, Krunic D, Mauer J, Lee YH, Granneman JG, Fischer L, Muller-Stich BP, Amri EZ, Kershaw EE, Heikenwalder M, Herzig S, Vegiopoulos A. Jak-TGFbeta cross-talk links transient adipose tissue inflammation to beige adipogenesis. Sci Signal 2018; 11(527).

Bayindir-Buchhalter I, Wolff G, Lerch S, Sijmonsma T, Schuster M, Gronych J, Billeter AT, Babaei R, Krunic D, Ketscher L, Spielmann N, Hrabe de Angelis M, Ruas JL, Muller-Stich BP, Heikenwalder M, Lichter P, Herzig S, Vegiopoulos A. Cited4 is a sex-biased mediator of the antidiabetic glitazone response in adipocyte progenitors. EMBO Mol Med 2018; 10(8).

Bayindir I, Babaeikelishomi R, Kocanova S, Sousa IS, Lerch S, Hardt O, Wild S, Bosio A, Bystricky K, Herzig S, Vegiopoulos A. Transcriptional Pathways in cPGI2-Induced Adipocyte Progenitor Activation for Browning. Front Endocrinol (Lausanne) 2015; 6: 129.

Betz MJ, Enerback S. Targeting thermogenesis in brown fat and muscle to treat obesity and metabolic disease. Nat Rev Endocrinol 2018; 14(2): 77–87.

Cannon B, Nedergaard J. Cultures of adipose precursor cells from brown adipose tissue and of clonal brown-adipocyte-like cell lines. Methods Mol Biol 2001; 155: 213–24.

Cannon B, Nedergaard J. Brown adipose tissue: function and physiological significance. Physiol Rev 2004; 84(1): 277–359.

Cherington V, Brown M, Paucha E, St Louis J, Spiegelman BM, Roberts TM. Separation of simian virus 40 large-T-antigen-transforming and origin-binding functions from the ability to block differentiation. Mol Cell Biol 1988; 8(3): 1380–4.

Chouchani ET, Kazak L, Spiegelman BM. New Advances in Adaptive Thermogenesis: UCP1 and Beyond. Cell Metab 2019; 29(1): 27–37.

Diaz MB, Herzig S, Vegiopoulos A. Thermogenic adipocytes: from cells to physiology and medicine. Metabolism 2014; 63(10): 1238–49.

Emont MP, Kim DI, Wu J. Development, activation, and therapeutic potential of thermogenic adipocytes. Biochim Biophys Acta Mol Cell Biol Lipids 2019; 1864(1): 13–9.

Forest C, Doglio A, Ricquier D, Ailhaud G. A preadipocyte clonal line from mouse brown adipose tissue. Short- and long-term responses to insulin and beta-adrenergics. Exp Cell Res 1987; 168(1): 218–32.

Hebert TL, Wu X, Yu G, Goh BC, Halvorsen YD, Wang Z, Moro C, Gimble JM. Culture effects of epidermal growth factor (EGF) and basic fibroblast growth factor (bFGF) on cryopreserved human adipose-derived stromal/stem cell proliferation and adipogenesis. J Tissue Eng Regen Med 2009; 3(7): 553–61.

Hirschberg V, Fromme T, Klingenspor M. Test systems to study the structure and function of uncoupling protein 1: a critical overview. Front Endocrinol (Lausanne) 2011; 2: 63.

Irie Y, Asano A, Canas X, Nikami H, Aizawa S, Saito M. Immortal brown adipocytes from p53-knockout mice: differentiation and expression of uncoupling proteins. Biochem Biophys Res Commun 1999; 255(2): 221–5.

Isidor MS, Winther S, Basse AL, Petersen MC, Cannon B, Nedergaard J, Hansen JB. An siRNA-based method for efficient silencing of gene expression in mature brown adipocytes. Adipocyte 2016; 5(2): 175–85.

Kazantzis M, Takahashi V, Hinkle J, Kota S, Zilberfarb V, Issad T, Abdelkarim M, Chouchane L, Strosberg AD. PAZ6 cells constitute a representative model for human brown pre-adipocytes. Front Endocrinol (Lausanne) 2012; 3: 13.

Klein J, Fasshauer M, Ito M, Lowell BB, Benito M, Kahn CR. beta(3)-adrenergic stimulation differentially inhibits insulin signaling and decreases insulin-induced glucose uptake in brown adipocytes. J Biol Chem 1999; 274(49): 34795–802.

Lee YH, Petkova AP, Konkar AA, Granneman JG. Cellular origins of cold-induced brown adipocytes in adult mice. FASEB J 2015; 29(1): 286–99.

Li Y, Fromme T, Schweizer S, Schottl T, Klingenspor M. Taking control over intracellular fatty acid levels is essential for the analysis of thermogenic function in cultured primary brown and brite/beige adipocytes. EMBO Rep 2014; 15(10): 1069–76.

Markussen LK, Isidor MS, Breining P, Andersen ES, Rasmussen NE, Petersen LI, Pedersen SB, Richelsen B, Hansen JB. Characterization of immortalized human brown and white pre-adipocyte cell models from a single donor. PLoS One 2017; 12(9): e0185624.

Marquez MP, Alencastro F, Madrigal A, Jimenez JL, Blanco G, Gureghian A, Keagy L, Lee C, Liu R, Tan L, Deignan K, Armstrong B, Zhao Y. The Role of Cellular Proliferation in Adipogenic Differentiation of Human Adipose Tissue-Derived Mesenchymal Stem Cells. Stem Cells Dev 2017; 26(21): 1578–95.

Moonen MPB, Nascimento EBM, van Marken Lichtenbelt WD. Human brown adipose tissue: Underestimated target in metabolic disease? Biochim Biophys Acta Mol Cell Biol Lipids 2019; 1864(1): 104–12.

Nedergaard J, Wang Y, Cannon B. Cell proliferation and apoptosis inhibition: essential processes for recruitment of the full thermogenic capacity of brown adipose tissue. Biochim Biophys Acta Mol Cell Biol Lipids 2019; 1864(1): 51–8.

Penfornis P, Viengchareun S, Le Menuet D, Cluzeaud F, Zennaro MC, Lombes M. The mineralocorticoid receptor mediates aldosterone-induced differentiation of T37i cells into brown adipocytes. Am J Physiol Endocrinol Metab 2000; 279(2): E386–94.

Rodeheffer MS, Birsoy K, Friedman JM. Identification of white adipocyte progenitor cells in vivo. Cell 2008; 135(2): 240–9.

Ross SR, Choy L, Graves RA, Fox N, Solevjeva V, Klaus S, Ricquier D, Spiegelman BM. Hibernoma formation in transgenic mice and isolation of a brown adipocyte cell line expressing the uncoupling protein gene. Proc Natl Acad Sci U S A 1992; 89(16): 7561–5.

Shinoda K, Luijten IH, Hasegawa Y, Hong H, Sonne SB, Kim M, Xue R, Chondronikola M, Cypess AM, Tseng YH, Nedergaard J, Sidossis LS, Kajimura S. Genetic and functional characterization of clonally derived adult human brown adipocytes. Nat Med 2015; 21(4): 389–94.

Sidossis L, Kajimura S. Brown and beige fat in humans: thermogenic adipocytes that control energy and glucose homeostasis. J Clin Invest 2015; 125(2): 478–86.

Wang W, Seale P. Control of brown and beige fat development. Nat Rev Mol Cell Biol 2016; 17(11): 691–702.

Widberg CH, Newell FS, Bachmann AW, Ramnoruth SN, Spelta MC, Whitehead JP, Hutley LJ, Prins JB. Fibroblast growth factor receptor 1 is a key regulator of early adipogenic events in human preadipocytes. Am J Physiol Endocrinol Metab 2009; 296(1): E121–31.

Wu J, Bostrom P, Sparks LM, Ye L, Choi JH, Giang AH, Khandekar M, Virtanen KA, Nuutila P, Schaart G, Huang K, Tu H, van Marken Lichtenbelt WD, Hoeks J, Enerback S, Schrauwen P, Spiegelman BM. Beige adipocytes are a distinct type of thermogenic fat cell in mouse and human. Cell 2012; 150(2): 366–76.

Xue R, Lynes MD, Dreyfuss JM, Shamsi F, Schulz TJ, Zhang H, Huang TL, Townsend KL, Li Y, Takahashi H, Weiner LS, White AP, Lynes MS, Rubin LL, Goodyear LJ, Cypess AM, Tseng YH. Clonal analyses and gene profiling identify genetic biomarkers of the thermogenic potential of human brown and white preadipocytes. Nat Med 2015; 21(7): 760–8.

Zilberfarb V, Pietri-Rouxel F, Jockers R, Krief S, Delouis C, Issad T, Strosberg AD. Human immortalized brown adipocytes express functional beta3-adrenoceptor coupled to lipolysis. J Cell Sci 1997; 110 (Pt 7): 801–7.

